# National-scale acoustic monitoring of avian biodiversity and migration

**DOI:** 10.1101/2024.05.21.595242

**Authors:** I. Avery Bick, Vegar Bakkestuen, Benjamin Cretois, Ben V. Hillier, John A. Kålås, Ingar J. Øien, Marius Pedersen, Kiran Raja, Carolyn M. Rosten, Marius Somveille, Bård G. Stokke, Julia Wiel, Sarab S. Sethi

## Abstract

Billions of birds migrate annually, triggered by endogenous behaviors but also by ecoclimatic drivers which are shifting with climate change. These dynamics play out over huge spatiotemporal scales, making monitoring of phenology challenging with traditional biodiversity survey approaches. In this study, over a complete spring migration season (April through June), we collected 37,429 hours of audio from 28 networked sensors in forests across Norway using a nationwide passive acoustic monitoring (PAM) system. We applied an open-source detection algorithm to automatically classify bird vocalizations; through expert validation we found the algorithm classified 57 species (14 full migrants) with at least 80% precision. Using these automated detections, we developed regional arrival curves for three common migratory passerines: Willow Warbler, Common Chiffchaff, and Spotted Flycatcher. We then demonstrate that PAM detections can be used to train audio species distribution models that map how species vocalization probability changes across Norway during spring migration. Lastly, we discuss how PAM can complement existing manual surveys to support the design and implementation of effective policy and conservation measures.

## Main

Around the world, climate change, habitat loss, and pollution are having severe impacts on biodiversity^1^, an integral aspect to each of the 17 United Nations’ Sustainable Development Goals (SDGs)^2^. Monitoring biodiversity is critical for understanding and mitigating the impact of such anthropogenic stressors, and for informing and evaluating conservation and land management strategies^3,4^. Historically, ecological studies relied on small teams or individual scientists, such as biologists studying the effects of pesticides on birds in the 1940s^5,6^. However, to expand the spatiotemporal scope of observations and enable estimates of population change, citizen science efforts began to emerge, with the North American Breeding Bird Survey in 1966 as a prime example^5^. Such efforts have since expanded greatly and are commonly used to track population change and inform conservation policy^7^. More recently, citizen scientist efforts have expanded to include the general public through apps such as eBird and iNaturalist, as well as Artsobservasjoner within Norway.

Breeding bird surveys are typically performed by knowledgeable volunteers^8^ and capture a snapshot of bird populations in the late-spring and early-summer across thousands of sites^8^. These surveys are a thorough and crucial means of estimating population trends and ranges for many species^7^. However, they are typically limited in their temporal coverage, as they are performed in specific temporal windows to survey breeding pairs and exclude passing avian migrants. This focus on population counts somewhat limits their effectiveness for studying migratory timing and phenology, such as shifts in Spring migratory bird arrival timing that are occurring with global warming^1,3^. Conversely, datasets which allow the public to record observations opportunistically (e.g., eBird) may provide wide spatial and temporal sampling coverage that can enable continental-scale studies of migratory phenology ^9^. Still, these datasets may suffer from temporal and spatial biases^10^ and a high variation in data reliability^11^. Ecologists have also long employed weather radars to track birds and migratory patterns over large distances^12^. While radar effectively tracks aggregate migrant behavior, such as identifying stopover locations^12^, it cannot easily distinguish among species, limiting its ability to study how short and long distance migrants are shifting their migration phenologies at different rates^3^, for example.

Other automated approaches are increasingly being used to complement manual surveys by improving the scale and resolution of biodiversity monitoring^13^. Passive acoustic monitoring (PAM) has shown particular promise due to the approach’s taxonomic breadth^14^ and scalability for both data analysis and collection^13^. Automated analysis of audio recordings is becoming commonplace, supported by the growth of powerful machine learning (ML) models^15^. ML algorithms such as BirdNET can accurately identify hundreds of species from recordings^16,17^, while others are used to detect biodiversity pressures such as noise pollution^18^, illegal logging^19^, or poaching^20,21^. Autonomous and real-time acoustic data collection via cellular network connection and self-powered units (via solar power or connection to a grid) is also growing^22,23^. PAM’s potential for continuous, accurate, and automated detection–in tandem with reduced labor requirements for monitoring and decreasing hardware costs^14^–promises to lower financial and manpower overheads for large-scale ecological monitoring. However, few studies^24^ have put data collection and analysis together to deliver biodiversity insight at scales and resolutions that would not be possible with traditional methods.

The study of avian species phenology is particularly well-suited for PAM. Many birds are highly vocal, and their dynamics and distributions extend over scales and resolutions that are not easily captured by traditional surveys. For example, at fine temporal resolutions, PAM can capture changes in bird vocalization behavior resulting from noise, such as passing snowmobiles^25^, with implications for land management policy. During longer-term biodiversity surveys, PAM enables continuous coverage, lowering manual labor requirements for repeat point count surveys, and increasing the probability of detecting rare species^14,26^. PAM is also particularly useful for studying passerine species during their critical Spring migration and during their breeding season, when they are particular vocal^27^. Indeed, avian vocalization rates often peak during vital activities such as pairing, nest building, and egg-laying and thus may be a strong measure of reproductive activity and success^27,28^. Furthermore, due to large existing vocalization libraries, off-the-shelf bird vocalization detection ML models cover thousands of species and are increasingly reliable^16,17,24^, reducing the need for training project-specific models. Overall, improved monitoring of avian phenology may build understanding around how stressors such as noise pollution^29^, habitat loss^30^, and climate change impact avian migration (e.g., a spring phenological mismatch between migratory arrival and greening^3,31^), informing conservation policy in an era of rapid ecological change^32^.

In this study, we perform one of the first national-scale acoustic surveys^33^ of avian phenology using autonomous acoustic recorders installed across almost the entire latitudinal extent of Norway with the goal of testing the applicability of PAM for large-scale monitoring of biodiversity and migration phenology. First, we compare automated vocalization detections against expert ornithologist validation to determine the precision of PAM detections in our study area. We then compare detections from PAM against those from the Norwegian Breeding Bird Survey and eBird within our study regions to determine how PAM may supplement these existing manual survey methodologies. Lastly, we test the applicability of PAM detections for training national-scale acoustic species distribution models (aSDMs)^34^. These aSDMs map the probability of detecting a species’ vocalizations based on ecoclimatic covariates and hold promise for tracking avian migration, mapping species detectability over time to inform effective manual surveys, and modeling species’ phenological responses to climate change^34^.

### PAM is reliable and fills data gaps in existing surveys

Our solar-powered, networked PAM stations were installed from Kristiansand on the southern tip of Norway, to Finnmark in the far north, approximately 1,700 km apart (Fig. 1a). Between April 1 and June 30, 2022, we collected 37,429 hours of audio data from 28 continuously-recording sites in eight regional clusters. Across recording stations, there was an there was an average 77% uptime (the percentage of time covered by recordings) from date of installation until the end of the study period. Having multiple recorders in each region provided data redundancy and covered for periods of power loss or recorder failure, such as with sites Kristiansand 1, Trondheim 3, and Bergen 3 (see SI Figure 1 for a detailed breakdown of recorder uptime). Recordings were saved locally on recorders in 5-minute files, then uploaded to a cloud server via mobile internet, where BirdNET was used to detect and classify bird vocalizations.

**Fig. 1.**
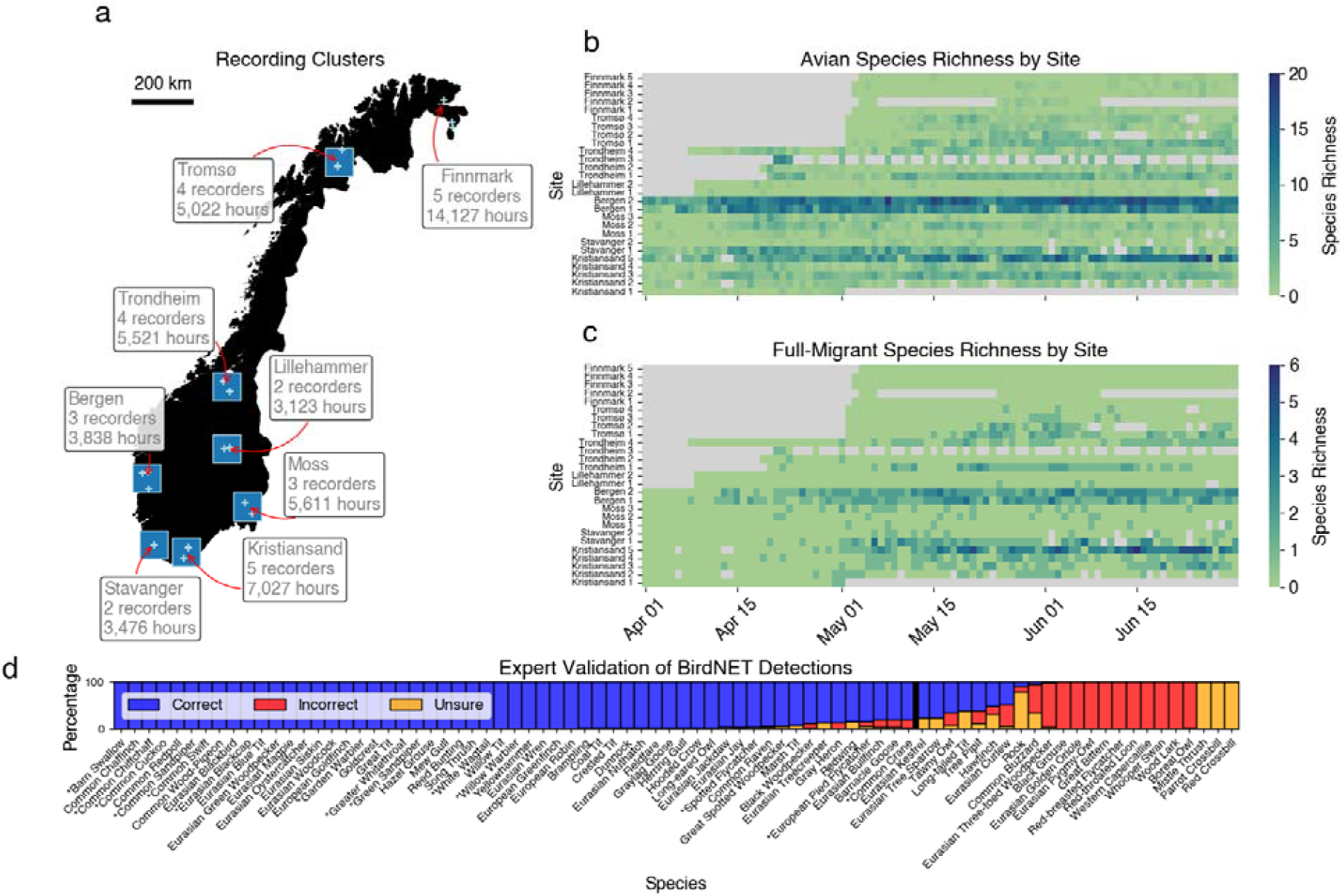
Passive acoustic monitoring captures avian species richness during migratory arrival and is highly precise in detecting dozens of species. **a**, 28 passive acoustic monitors deployed across Norway in 2022 are grouped into eight regions. Blue boxes (90 km x 90 km) were drawn centered on the geometric centroid of each region’s recorders; these boxes are used to aggregate manual bird survey data for comparison to audio detections, and to spatially average daily NDVI data^36^. No box was drawn for the Finnmark cluster, as no detections were above our model confidence threshold in this region. Light blue crosses represent coordinates of recorders. Within regions, some recorders were placed close to each other, but maintained acoustic independence (i.e., >100m apart, see Sethi et al 2020^37^). **b**, Species richness of 57 avian species and **c**, 14 full-migrant species vary across Norway temporally and latitudinally. Passive acoustic monitoring can provide near-continuous monitoring over long periods, but redundancy is key as equipment in the field may fail (such as site Kristiansand 1 in early May) or solar panels may receive insufficient sunlight. Days with no recorder uptime are shown as gray. **d**, Expert ornithologist validation of BirdNET species identification performance for all 67 species with _≥_50 detections over _≥_0.80 model confidence. Of 67 species, 57 species are detected with _≥_80% precision across 50 randomly sampled detections, indicated by species to the left of the thick black line. Full migrant species are identified by a “*” before their common name.

We utilized an expert validation methodology to understand the per-species precision of BirdNET classifications in our study area and to focus our analysis on species that the model classified with the highest precision. Fifty randomly selected BirdNET detections, and their corresponding audio, were provided to an expert ornithologist for each of 67 species which had at least 50 BirdNET detections with a model confidence^35^ over 0.80. After expert labeling of these recordings, we found that BirdNET correctly identified 57 species with greater than 80% precision and 30 of these with 100% precision (Fig. 1d). Of these species, 14 were full migrants which are not present in Norway during the winter. For these migrants, we observed increasing species richness through the spring months as well as latitudinal differences in biodiversity and migratory arrival timing (Fig. 1c).

We then focused further on three common migratory passerines detected with 100% precision (Fig. 1d): Willow Warbler (WW) (*Phylloscopus trochilus*), Common Chiffchaff (CC) (*Phylloscopus collybita*), and Spotted Flycatcher (SF) (*Muscicapa striata*)), which had sufficient high-confidence detections (≥ 20) to calculate 5% detection threshold arrival dates across several sites. We compared the survey temporal coverage, total detections, and sensitivity of PAM with two comprehensive avian occurrence datasets available for Norway: opportunistic encounters recorded on eBird^38^, and standardized line transect point counts from Norsk Hekkefuglovervåking, also known as the Norwegian Breeding Bird Monitoring Scheme (BBS)^8^, which is performed in the breeding season between late May to early July, after which most migrant species have arrived. PAM had greater total species observations than manual surveys for WW across the study period within our study regions (N_PAM_ = 687, N_eBird_ = 102, N_BBS_ = 135), CC (N_PAM_ = 300, N_eBird_ = 48, N_BBS_ = 63), and SF (N_PAM_ = 192, N_eBird_ = 60, N_BBS_ = 45), primarily driven by the large number of days when acoustic data was recorded but no eBird or BBS survey were performed (light blue, Fig. 2). We note a distinct two-peak pattern of vocalization frequency over time for the three species (Fig. 2a,b,c), that has previously been observed for Willow Warbler^28,39^ and other passerines^28,40^. The first peak in mid-May may correspond with the pre-breeding season (Fig. 2a), with males establishing territory and performing courtship^39^. This is followed by a drop in vocalization during pairing, then a second increase during laying and incubation, which may be related to courtship for second clutches^28,39^. However, the vocalization of migrating birds stopping over in Bergen may also contribute to this two-peak pattern.

**Fig. 2.**
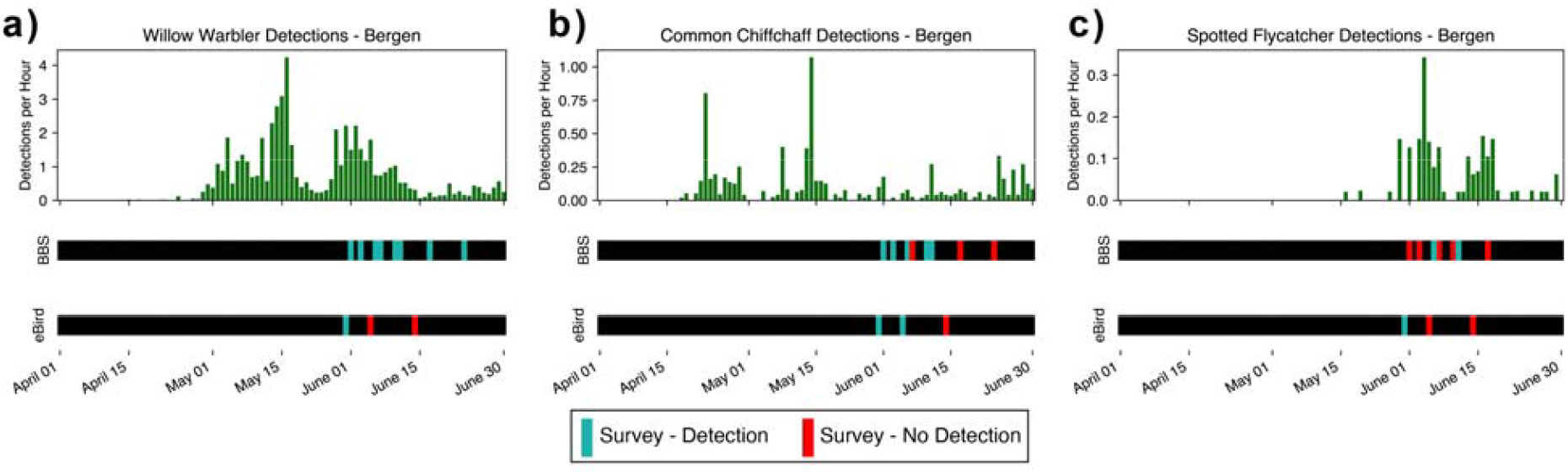
Comparison of data acquired through passive acoustic monitoring (PAM), standardized point-count (BBS), and open citizen science surveys (eBird) for detecting three species of migrating passerines. In Bergen, the PAM recorders capture species vocalization frequency (green bars) across the spring migration period for: a) Willow Warbler, b) Common Chiffchaff, and c) Spotted Flycatcher. The data from BBS and eBird also confirm the presence of these species through detections (teal lines) and non-detections (red lines).

As BBS is only performed during a limited time window, we also compared detections solely on days where both BBS and PAM data were available. BBS generally had a higher probability of detecting these species on days when PAM did not; there were 42 days where BBS detected WW, CC, and SF and PAM did not. Conversely, there were only 13 days where only PAM detected these species. eBird also exhibited a higher detection sensitivity than PAM on days where both surveys were conducted, with 21 to 14 unique daily detections, respectively. These results highlight the synergy between these distinct survey methods. PAM provides greater sampling effort temporally, giving phenological context for manual surveys which are concentrated temporally during breeding season. In contrast, BBS had a greater spatial sampling effort when compared to the two to five PAM recorders per 90 km x 90 km region. However, increasing the number of PAM recorders in a given region would improve spatial sampling and likely increase the detectability of these species^41^.

### PAM enables avian phenology monitoring at a national scale

We set a threshold of 5% of cumulative detections^42^ across our study period to determine a species’ arrival. In Bergen, where we have sufficient (≥ 20) high confidence detections to determine arrival date for each migratory species, we estimated that CC, WW, and SF arrived on April 23, May 4, and May 27, respectively (Fig. 3c, Methods). Indeed, CC winters further north than WW and SF^43–45^, which are long-distance migrants, and thus CC generally arrives earlier^46^. The timings also align with these species’ respective food sources. WW^43^ and CC^44^ primarily eat crawling insects, larvae, and berries while SF^45^ eat flying insects, for which peak populations occur later in the Spring season in northern regions^47,48^. The migratory arrival order seen in Bergen is consistent for other regions, with WW arriving before SF in Kristiansand, and CC arriving before WW in Trondheim (Fig. 3b). Based on the arrival times in southernmost region Kristiansand and northernmost region Tromsø and the distance between the region centroids, WW traveled northward at approximately 67 km/day, comparable to previous measurements of WW southward autumn migratory rate of 40.6-80.5 km/day using ringed birds^49^, though daily migration rates differ between spring and autumn migration^50^.

**Fig. 3.**
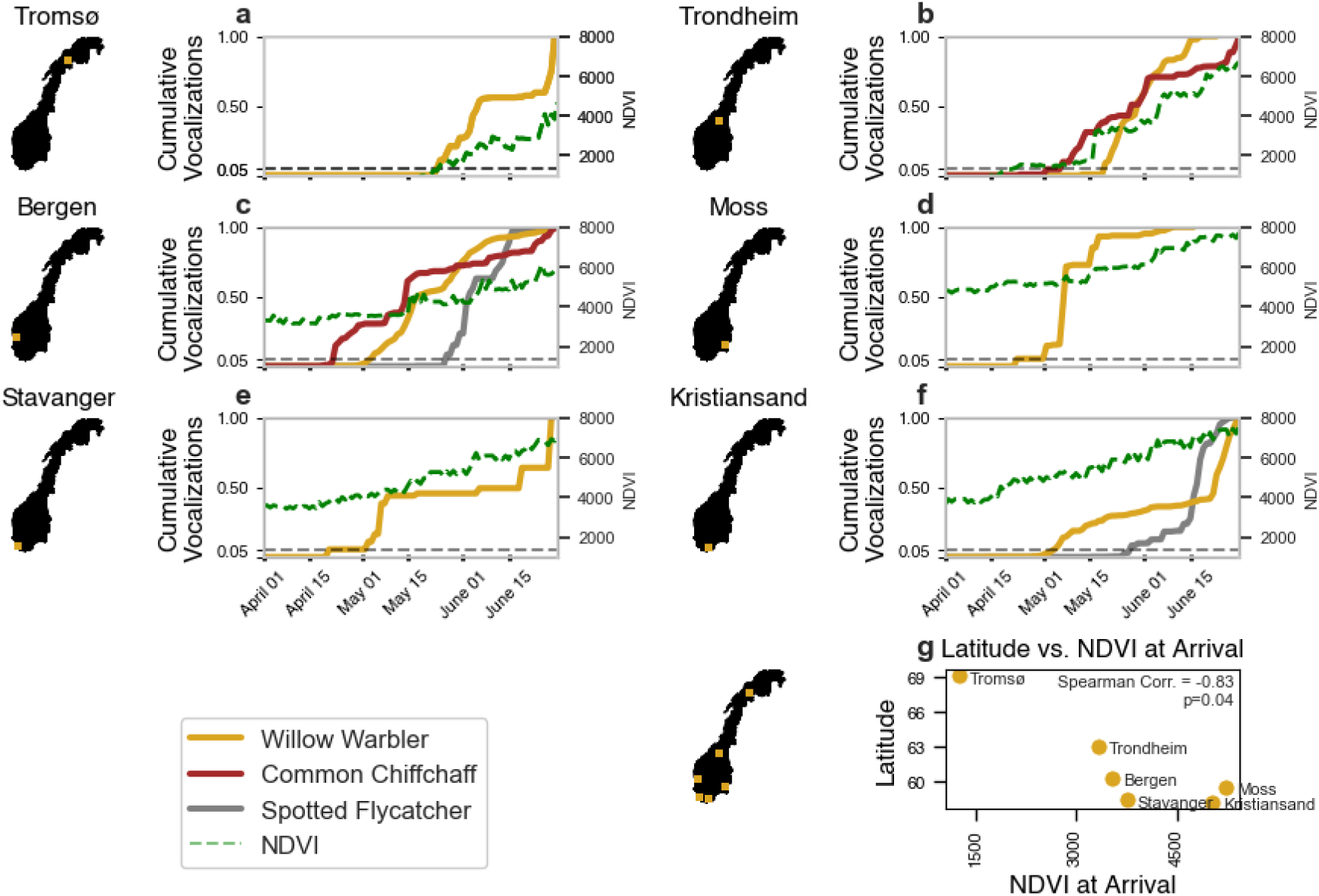
Cumulative audio detections of migratory passerines demonstrate interspecies and large-scale variation in migration timing phenology across Norway. **a-f**, Passive acoustic monitoring shows ecologically consistent species arrival patterns across Norway^36^. Willow Warbler and Common Chiffchaff, whose diets include primarily larvae and crawling insects, arrive earlier than Spotted Flycatcher, which have a diet of flying insects that emerge later in the season. Cumulative detections are shown only for species with at least 20 high-confidence detections, sufficient for calculating migratory arrival based on a 5% threshold. **g**, Willow Warbler arriving at higher latitudes arrive at a much lower greenness index than those at lower latitudes (Spearman _ρ_=-0.83, p=0.04, N=6).

We found that WW generally arrives at a lower greenness at higher latitudes, as measured by satellite-derived Normalized Difference Vegetation Index (NDVI)^51^ (Spearman ρ=-0.83, p=0.04). The lower NDVI at arrival in higher latitudes highlights the delicate balance that migratory species must strike regarding their arrival timings to spring breeding grounds. Migrating birds breeding in arctic and sub-arctic regions contend with short breeding seasons, so they carefully balance their arrival times to match the timing of snowmelt and greening of northern birch forests^52^, ideally aligning peak food abundance to when chicks are present in their nests^46,53^. Mistiming migration to these colder, northern areas may lead to mortality^46,54^ and climatic changes in wintering areas in Africa seem to have driven WW to arrive approximately half a day earlier to the Tromsø region each year since 1995^46^, potentially threatening this relationship between evolved arrival timing and food availability. Longer-term PAM installations in arctic and sub-arctic regions^32^ could aid in tracking migratory arrival times and phenological mismatches with spring greening, due to PAM’s ability to operate continuously in remote regions which may have limited manual survey data.

We next hyperparameter-tuned and trained random forest regression models^55^ to act as aSDMs^34^, predicting the weekly probability of detecting WW and SF vocalizations based on the values of several ecoclimatic features: weekly averages of NDVI^51^, maximum^56^, minimum^57^, and mean^58^ temperature, and precipitation^59^, and recorder elevation^60^. The aSDMs allowed us to extrapolate spatially beyond our sampling sites to other forested regions^61^ similar to where we sampled, mapping migratory waves of birds based on their probability of vocalization (Fig. 4) at a resolution of 0.2 degrees, approximately 10 km.

**Fig. 4.**
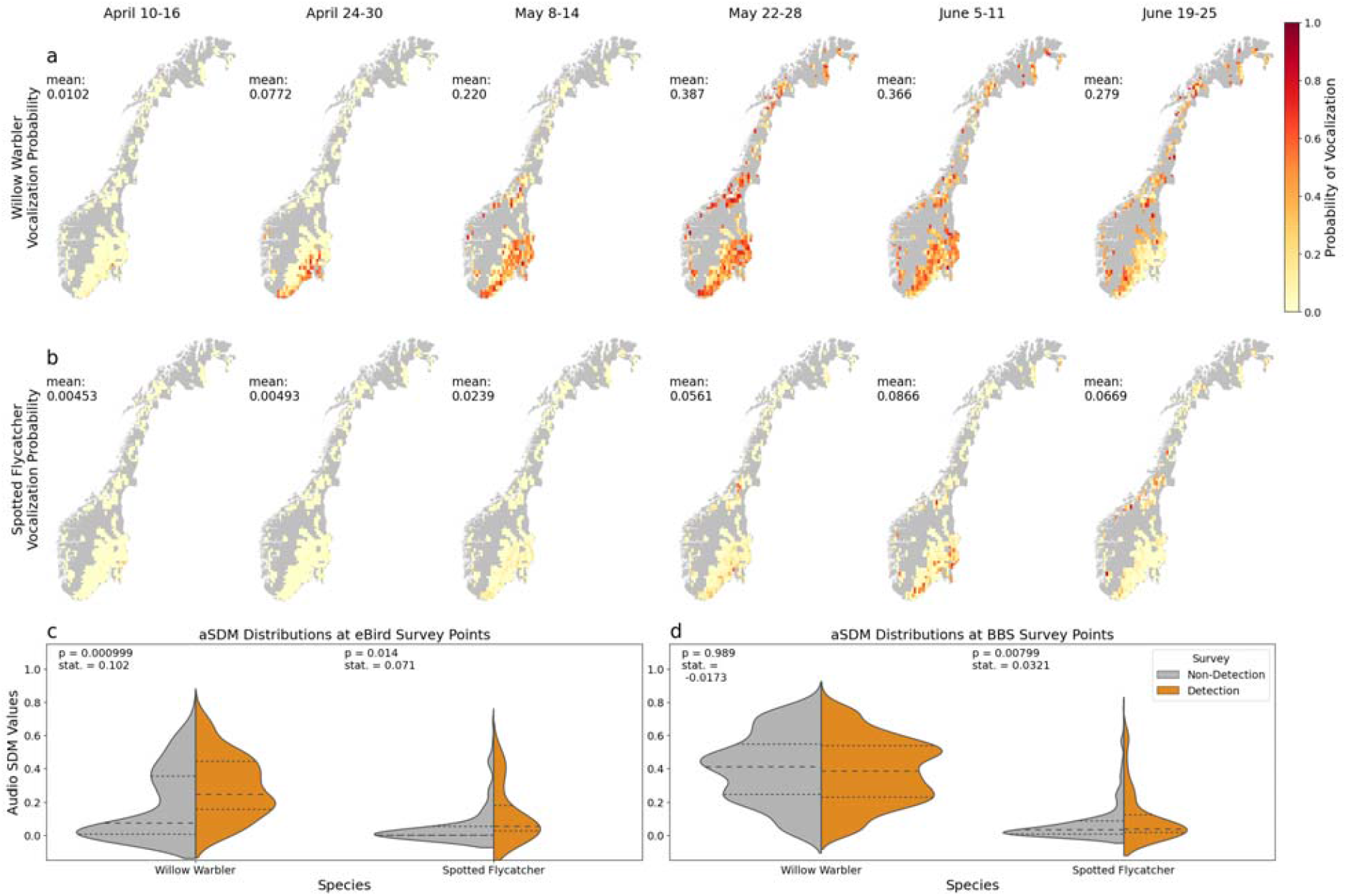
We trained a simple machine learning model to predict the probability of hearing a given species’ vocalization based on several ecoclimatic variables. These models are known as audio species distribution models (aSDMs). Here, the vocalization probability output can be interpreted as an analogue of migratory species detectability across Norway. Weekly vocalization probability in forested areas across Norway^36^ for **a**, Willow Warbler and **b**, Spotted Flycatcher shows spatiotemporal species arrival patterns and captures later arrival of Spotted Flycatcher relative to Willow Warbler. **c**, At eBird survey points, the distribution of aSDM values is significantly greater when the surveyor detected WW and SF versus when the eBird surveyor did not detect the species. **d**, For BBS survey points, the distribution of aSDM values is significantly greater when the surveyor detected SF, but not for WW. Dashed lines represent distribution medians and dotted lines represent quartiles.

With the national scale aSDMs, we observed that WW’s earlier arrival and at a higher relative abundance than SF across Norway, as demonstrated by their vocalization probabilities (Fig. 4a,b). Between May 8-14, the model predicts that Willow Warbler are largely not yet vocalizing in the northern Tromsø region, but that these areas are populated two weeks later (May 22-28). This aligns with both the raw species detections (Fig. 3a) and the estimate of May 14 as the first arrival date of Willow Warbler in Troms County by Barrett^52^, though this study was performed in 2002 and first arrival times may have since advanced^46^.

To test the agreement of the aSDMs with eBird and BBS survey data in forested regions, we sampled the distributions of aSDM values at survey observation points, observing whether higher aSDM vocalization probabilities were associated with detections from manual surveys (Fig. 4c,d). The distribution of aSDM values at the locations of eBird detections were higher than aSDM values at locations of non-detections by permutation test for WW (p≤0.001, t=0.102, N_detections_=146, N_non-detections_=226, μ_detections_=0.292, μ_non-detections_=0.190) and SF (p=0.014, t=0.071, N_detections_=20, N_non-detections_=352, μ_detections_=0.108, μ_non-detections_=0.037) (Fig. 4c,d). Similarly, Bota et al 2020 found that the vocal activity rate of European Bee-eaters, as detected by PAM recorders, correlated positively with their inclusion in complete checklists from citizen scientists^62^. However, we see that the aSDMs for WW do not significantly agree with the BBS observations (p=0.97, t=-0.016, N_detections_=2080, N_non-detections_=791, μ_detections_=0.379, μ_non-detections_=0.395) (Fig. 4d), perhaps due to the limited temporal range of BBS in late May and June. WW are nearly ubiquitous across Norwegian forests in this time period (Fig. 4a); this may reduce the ability of the aSDM model to identify relevant ecoclimatic features for predicting vocalization probability. In contrast, the BBS observations agree with the aSDM for SF (p=0.00799, t=0.0300, N_detections_=97, N_non-detections_=2774, μ_detections_=0.104, μ_non-detections_=0.0704), which have a more heterogenous distribution (Fig. 4b).

## Discussion and Future Work

Our results show that national-scale PAM deployments can deliver a step change in both the resolution and scale at which we can track avian phenology when compared to existing manual surveys. Pairing networked sensors and automated analyses with one-off expert validation allowed us to track phenology reliably and in near real-time, and required minimal cost and effort to deploy and maintain. For example, from PAM detections we calculated arrival times of several migrating passerines which align with their ecological niches and results from previous studies^46^. We also found a distinct two-peak pattern in vocalization frequency over the Spring consistent with previous work by manual surveys^28^.

However, even with improved temporal resolution and lower labor requirements, PAM methods should be viewed as a complement to existing manual observation methods, such as breeding bird surveys and eBird. Each of these survey types have a distinct method and threshold for determining a species detection. Thus, PAM detections may fill gaps in manual surveys and vice-versa. For example, BBS requires both the male and female in a breeding pair to be observed and thus offers a rigorous estimation of a species’ breeding population at each survey point. However, the manual labor required and limited time frame restrict its temporal resolution. PAM typically does not distinguish between male or female individuals and may struggle to detect less-vocal species. PAM does offer much higher temporal resolution than BBS, which enables estimation of a species’ migratory arrival dynamics in a given area and study of its vocal phenology. The vocalization probabilities captured by PAM-derived aSDMs also offer a metric of species detectability^11,63^ which may inform the timing of both breeding bird surveys and more casual citizen scientists by indicating the annual and diurnal periods of maximum detectability for vocalizing bird species. As aSDM models function by relating ecoclimatic features to vocalization probabilities, they may be extended to project how conditions suitable for species vocalization may change over time with climate change. Desjonquères et al 2022, for example, predicted how the spatial vocalization probability of the Iberian Tree Frog may change with future shifts in temperature and precipitation according to the Representative Concentration Pathway 8.5 climate projection^34^.

Unlike breeding bird surveys, which are typically performed after Spring arrival for all migratory species, PAM cannot easily distinguish between vocalizations of resident birds in an area versus those who are passing migrant individuals. Similarly, during autumn migration, distinction cannot be easily made between adult and hatch-year individuals. PAM also does not typically discern whether detections are from individual birds or multiple, so a single individual may account for many of the detections at a given recorder. However, progress is being made on this front^64^, with the use of multiple microphones in a single recording array enabling spatial triangulation of vocalizing individuals^65^ and the use of specialized detection models that can distinguish subtle differences in song or call between individuals^66^. Such advancements in individual differentiation could improve PAM-derived population density estimates and supplement population counts from breeding bird surveys. The general avian vocal detection algorithm used here, BirdNET, also does not classify species detections as a bird call versus a song, though it is trained on examples of both^17^. However, custom models can be trained to distinguish between a species’ call and song^67^.

The expert validation approach employed here enabled estimation of BirdNET model precision, or the fraction of BirdNET detections for a given species with correct classifications. However, this approach is limited by inability to calculate recall, the fraction of true detections for a given species that were captured by BirdNET. Calculating recall would require each BirdNET detection to be manually checked for all species, a challenge considering there were 33,643 detections with a model confidence over 0.80. We argue that precision provides a more useful metric, critical for ensuring reliable detections in remote areas with lower bird populations and for facilitating stakeholder trust in model results. However, by thresholding detections with a model confidence over 0.80 for validation, we subsequently filtered out 81% of the total detections from our recorders. This limited the number of detections available to train aSDM models and reduced the sample size of detections for determining species’ arrival curves. This was especially relevant the recording cluster in Finnmark, which had several Willow Warbler detections, but none above the 0.80 model confidence threshold. Depending on goals, future studies may consider lowering thresholds for model confidence, or setting model confidence thresholds on a species-by-species basis^16^. Financial and practical limitations also affected the number of recorders deployed to each region, making our study sensitive to failure of some recording devices (see Fig. 1) and introducing possible spatial bias in the total number of detections by region. However, when calculating PAM detections per hour (Fig. 2a,b,c), we partially corrected for this bias by normalizing regional daily detections by the total regional daily number of hours recorded.

Future studies may take advantage of the fact that sensors are becoming more affordable and more reliable^20^ while machine learning models for identifying species by vocalization are improving in accuracy, taxonomic breadth (e.g., insects, mammals), and computational efficiency^15^, making large-scale PAM surveys more readily implementable. In fact, the TABMON^68^ project has already collected data from a continental-scale real-time PAM system built along avian migratory pathways across Europe. As with the Dutch ARISE program^69^, PAM can also be integrated with other sources of species observations, such as camera traps^70^, drones^71^, citizen science^72^, and formal surveys^8^ to provide more comprehensive information on biodiversity and ecological health. Overall, we show here that PAM facilitates ecological research and monitoring on large spatial scales and fine temporal scales, supporting manual survey and population count efforts and offering a higher resolution understanding of animal phenology. A synergy of automated techniques and manual surveys can pave the way for effective data-driven conservation and policy decision making, leading to improved outcomes for biodiversity globally.

## Materials and Methods

### Recording Site Selection

Recording sites were located at BBS survey points (within 100m) to maximize compatibility between the two datasets. The sites were also restricted to forested areas, as determined by the “Forest” IPCC Class in 300m resolution land cover data from Copernicus ^61^. When possible, sites were chosen within 200m of accessible roads^73^ or trails for ease of deployment and within areas with 4G data coverage. To maintain confidentiality of BBS survey points, random noise of up to 1 km was added to the location of each recorder in the publicly available version of this dataset.

Across Spring 2022 in Norway, 28 recording stations were installed across eight regions (from south to north): Kristiansand (5), Stavanger (2), Moss (3), Bergen (3), Lillehammer (2), Trondheim (4), Tromsø (4), and Finnmark (5). The different numbers of recorders per region were due to physical and logistical difficulties installing records in certain locations, including land access or insufficient sunlight to use the solar panel . While uncorrected spatial biases could exist in our analyses due to variable numbers of recorders per region, we do correct for total number of hours recorded per region to partially alleviative this, as described below in the description of methods for estimating species arrival.

These recorders were then grouped into regions, defined by a 90 x 90 km box based on the centroid of the recorders. These dimensions represented the largest size possible without overlap between regions. The Finnmark cluster of recorders was dropped from species arrival analysis, as only one valid detection of a species of interest for this study was found. However, detections from these sites are included in Figure 1 and for training and evaluating the audio species distribution models (Fig. 4). Sites within clusters were spread such that each could be considered acoustically independent (i.e., >100m apart, see Sethi et al 2020^37^).

### Audio Collection

In total, 73,516 hours of audio was collected from March to October 2022. Audio at each site was collected using a Bugg device 1-2m from ground level. Recorders operated continuously, though devices were powered through using 50-100W solar panels and 20-30 Ah 12V batteries and thus were sensitive to shade and cloud cover. Still, recorders had an average of 77% uptime from their installation until the end of the study period (see SI Fig. 1). The Bugg recorders used a built-in Knowles SPH0641LU4H-1 omnidirectional PDM MEMS microphone, with a 100Hz-80KHz dynamic range, which was weatherproofed using an acoustic ePTFE membrane ^74^.

Files were saved locally in five-minute increments at a 44.1 kHz sampling rate in compressed MP3 format (VBR0 using FFMPEG’s LAME codec), then uploaded to a Google Cloud server via the 4G network. Previous work has found MP3 compression at such high quality has little effect on downstream eco-acoustic analyses ^75^. Where network service was unavailable, audio data was manually uploaded upon device retrieval in Fall 2022.

### Species Detection

Audio data was processed on a custom Google Cloud web app using BirdNET-lite^76^ to extract and identify vocalizations to species level in near real-time^17^. In total, 176 unique species were identified in this dataset by BirdNET-lite. Of these, 67 species had at least 50 detections with a model confidence over 0.8. For this subset, we provided an expert ornithologist with 50 random detections for each species, sampled across all sites without stratification, with each species taking approximately 20-30 minutes to validate. Of these, the expert determined that BirdNET-lite correctly identified 57 species with 80% or greater precision. Of these species, 14 are full migrants to Norway each Spring. Expert validation notes are available in the online Zenodo repository for this paper.

### Species Arrival

We calculated species arrival for a subset of three fully migrant passerines that were commonly detected in our dataset and had high detection accuracy from BirdNET: Willow Warbler (*Phylloscopus trochilus*), Spotted Flycatcher (*Muscicapa striata*), and Common Chiffchaff (*Phylloscopus collybita*). We defined the study period as April 1 through June 30, 2022, by which most migratory bird species have arrived across Norway. In the Northern regions of Finnmark and Tromsø, recorders were installed in early May. With snow still covering the area, this temporal discrepancy is unlikely to miss observations–the first Willow Warbler in Tromsø was detected in mid-May, agreeing with previous manual estimates of first arrival^52^.

We estimated the date of migratory arrival for each species at a given region by setting a threshold of 5% total detections across our study period, a common method for setting arrival dates in migratory studies^42^. We chose this method over others, such as establishing migratory arrival by the first or second species detection, due to the potential for stopover migrants^46^ biasing arrival dates as they fly to northern nesting grounds. Due to our use of a 5% threshold, when calculating arrival for a given species, we only considered regions with more than 20 valid detections. To correct for varying spatiotemporal sampling effort, the daily detections of each species were normalized by the daily sum of recorder uptime within each region. To calculate the percentage of total vocalizations over time, we first took a cumulative sum of these normalized uptimes across the study period, then normalized these from 0 to 1. We estimated the date of species arrival as the first day with ≥ 5% of total vocalizations. We then plotted arrival curves along with normalized difference vegetation index (NDVI), sourced from the daily 0.05 degree resolution gap-filled MODIS Vegetation Index dataset^51^. Daily NDVI per region was calculated by averaging all NDVI cells within each 90×90 km region. We used Spearman rank correlation to measure the monotonic relationship between regional NDVI values on WW arrival days and the latitude of the region centroids.

### Comparison of Audio to Manual Survey Methods

#### eBird and BBS Surveys

Raw eBird surveys^38^ were aggregated within each 90x90 km region, then filtered by several criteria to standardize data^11^. First, to ensure that pseudo-absences could be assumed from eBird surveys, only complete checklists were included, those where observers marked that all observed species were included. Surveys with high effort–those traveling over 5km or lasting longer than 90 minutes–were dropped to ensure that observations occurred close to the provided survey start coordinates^11^. Duplicate surveys that appear in group surveys were dropped. Lastly, surveys in non-forested areas, as identified by the Copernicus data set^61^, were dropped to ensure comparability against the audio dataset. Lastly, surveys were filtered to the elevation range of our recording stations, 73m to 764m.

Typically, Norwegian Breeding Bird Survey (BBS) data is spatially aggregated to protect the specific coordinates of survey points from increased traffic which could affect avian nesting habits. Non-public BBS data was obtained from the survey organizers^8^ for the three migratory passerines of interest. The dataset consists of observations of the number of breeding pairs of each species at 9,257 unique points across Norway by experienced volunteers. Many volunteers are trained by attending the BIRD ID course at Nord University, but a prerequisite for all is good knowledge of bird sounds and appearance. Regional coordinators communicate with observers before they perform surveys, and observations are skipped in rare cases where poor knowledge or reduced hearing ability is suspected by the survey organizers. The BBS is carried out from 20 May through 10 July each year, after all species have arrived at their breeding sites, to be able to exclude individuals that are traveling. Volunteers travel along transects, making observations at several points. Between 4AM and 10AM, they count the number of breeding pairs they observe in 5 minutes at each point, visiting each point only once^8^. These observations were also filtered to forested regions and to the elevation range (73m to 764m) of our recording stations.

#### Audio Species Distribution Models

We trained and cross-validated a random forest regression model on several weekly ecoclimatic covariates^77^ to predict the probability of vocalization^34^ for Willow Warbler and Spotted Flycatcher. The weekly binary detection (0,1) of a species at each of the 28 sites served as the target variable, with weekly averages of NDVI^51^, maximum^56^, minimum^57^, and mean^58^ temperature, and precipitation^59^, as well as site elevation^60^ as the predictor variables. Predictor variables were extracted for each site’s coordinates. In Finnmark and Tromsø, where snow cover continues late into the spring, several weeks of NDVI coverage were not available from the MODIS dataset^51^ and were zero-filled. The random forest regression models were hyperparameter-tuned using a random grid search approach over 1,000 iterations. The model parameters with the best mean squared error performance on the test set were chosen via 4-fold cross-validation. The tuned model parameters and train and test set errors are available in SI Table 4.

We then generated weekly aSDMs using the trained models for each species. Covariate rasters were resampled to a common resolution of 0.2 degrees (epsg:4326) (approximately 10 km) and used to predict probability of vocalization across Norway. To restrict our vocalization probability maps to ecosystems similar to our study sites, we employ a masking step to filter out non-majority-forested cells from the aSDM based on the 300m resolution Copernicus land cover dataset^61^. Lastly, aSDMs were then compared against the filtered complete eBird and BBS lists (i.e., presence and absence included). For each month, we compared the distributions of aSDM values with positive survey detections (1) and for the survey absences (0) using a permutation test with 1,000 resamples and alternative “greater”. This test evaluated whether the means of the distributions of aSDM values were greater when eBird or BBS detects a species, versus when they do not.

## Acknowledgements

We thank Tom Roger Østerås of Norsk hekkefuglovervåking for his expert opinion in validating BirdNET species detections, NINA field staff for assistance with audio recorder setup, and the Norwegian Environment Agency (Miljødirektoratet) for funding of data collection. We also thank the homeowners who allowed us to install acoustic recorders on their property for this project. IAB was funded by a doctoral research grant 323294 from the Research Council of Norway.

## Author Contributions

CR, SSS, and BC organized the Sound of Norway (thesoundofnorway.com) project which facilitated this study. IAB identified suitable locations for acoustic recorders and worked with CR to receive landowner permissions to deploy recorders. CR, SSS, IAB, and JW deployed and collected acoustic recorders. IAB, SSS, MS, and BH developed methodologies and research objectives, with advisement from VB, MP, and KR. IAB performed all data processing and modeling and generated all figures. IAB drafted the manuscript with significant contributions from SSS. All coauthors reviewed and approved the final manuscript.

## Data Availability

All code, manual survey data, detection validation notes, acoustic detections, and ecoclimatic covariates required to generate each figure in this study are available at doi.org/10.5281/zenodo.11238950. Random noise of up to 0.01 degrees has been added to all coordinates of recording stations and Norwegian Breeding Bird survey points to maintain the confidentiality and data integrity of the survey.

## Supporting Information

**SI Figure S1.**
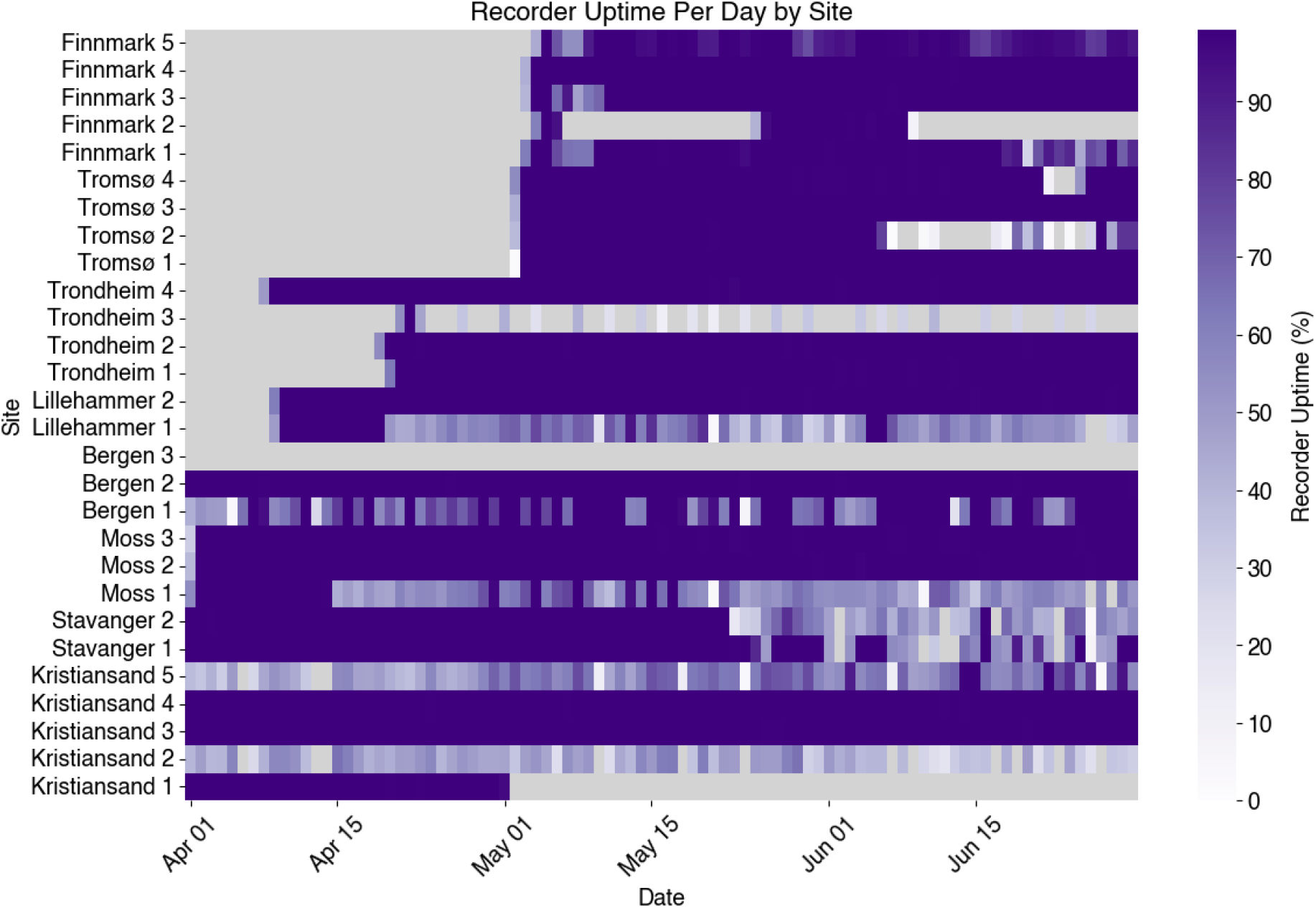
SI Acoustic recorder uptime percent for each site across the study period. Grey areas indicate no data availability.

**SI Table 1.** Total detections of Willow Warbler (WW), Spotted Flycatcher (SF), and Common Chiffchaff (CC) by study region and monitoring method.

| Region | Species | BBS Detections | eBird Detections | Audio Detections |
| --- | --- | --- | --- | --- |
| Kristiansand | WW | 95 | 34 | 425 |
| Bergen | WW | 65 | 1 | 2431 |
| Trondheim | WW | 88 | 5 | 158 |
| Tromsø | WW | 36 | 1 | 124 |
| Stavanger | WW | 48 | 2 | 29 |
| Moss | WW | 88 | 18 | 41 |
| Lillehammer | WW | 58 | 0 | 3 |
| Kristiansand | SF | 4 | 4 | 97 |
| Bergen | SF | 2 | 2 | 88 |
| Trondheim | SF | 3 | 0 | 4 |
| Tromsø | SF | 0 | 0 | 0 |
| Stavanger | SF | 0 | 0 | 8 |
| Moss | SF | 13 | 10 | 1 |
| Lillehammer | SF | 0 | 0 | 0 |
| Kristiansand | CC | 9 | 2 | 1 |
| Bergen | CC | 8 | 2 | 339 |
| Trondheim | CC | 51 | 9 | 151 |
| Tromsø | CC | 0 | 5 | 0 |
| Stavanger | CC | 2 | 2 | 0 |
| Moss | CC | 6 | 7 | 0 |
| Lillehammer | CC | 3 | 0 | 0 |

**SI Table 2.**
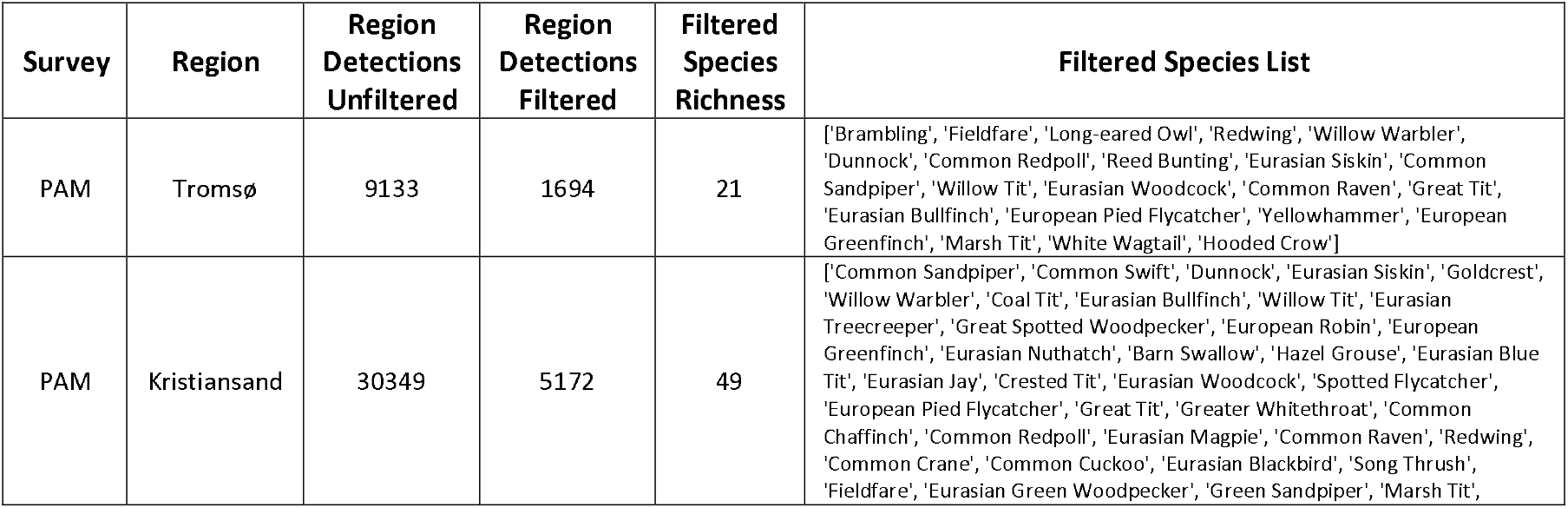
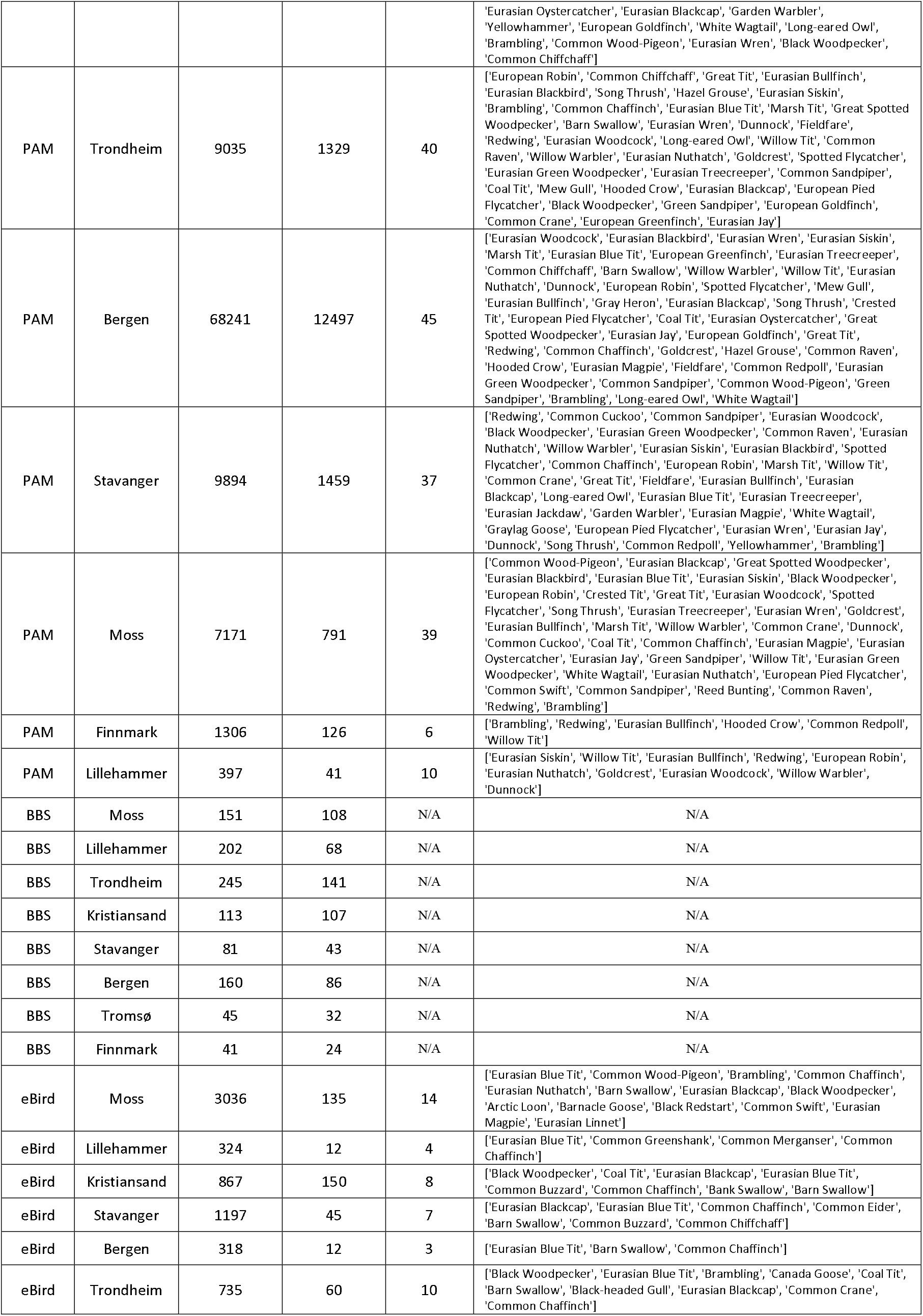
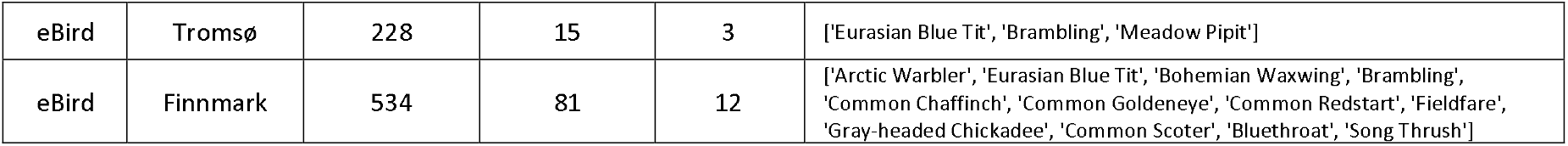
Regional total bird detections for three monitoring methods (PAM, BBS, and eBird) for April through June 2022. Detections are shown before and after data filtering steps. Species richness information is shown for filtered detections. For PAM, the “region detections unfiltered” column represents the total number audio detections per region for 57 species for which BirdNET was shown to have at least 80% precision; the “region detections filtered” column represents detections filtered for high model confidence (≥0.80). For BBS, the “region detections unfiltered” column represents all detections for WW, SF, and CC only within each region, as only data for these species were provided by BBS. For BBS, the “region detections filtered” column was additionally filtered to detections within the altitude range of our recorders and for detections in forested areas. For eBird, the “region detections unfiltered” column shows total species detections for all species after dropping any duplicates stemming from group surveys. For eBird, the “region detections filtered” column was further filtered to remove high effort surveys, surveys not within forested regions, non-complete surveys, and surveys not in the altitude range of recorders. See the Methods section for more details on filtering. The “filtered species richness” and “filtered species list” columns account for species present in the filtered data.

**SI Table 3:**
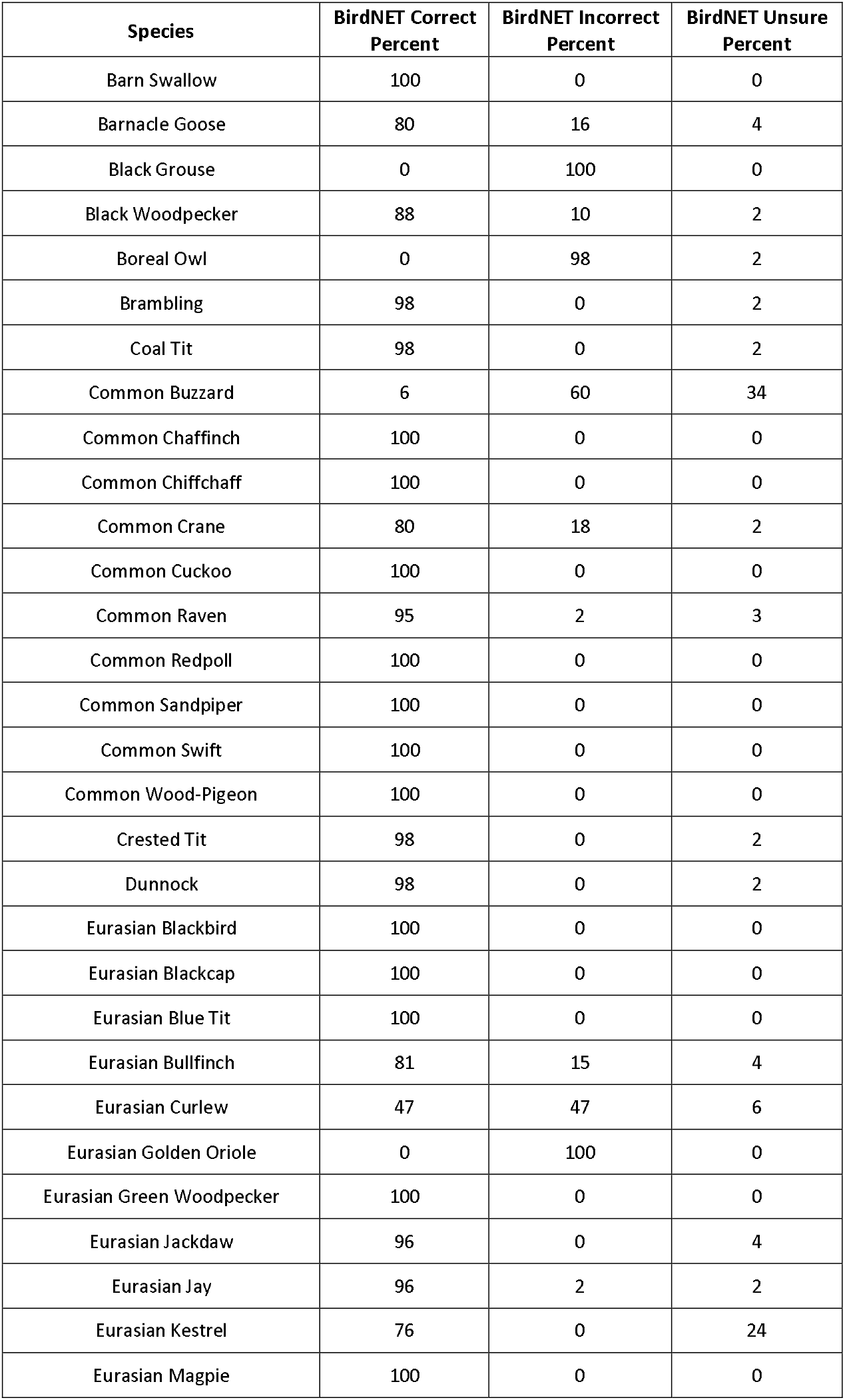
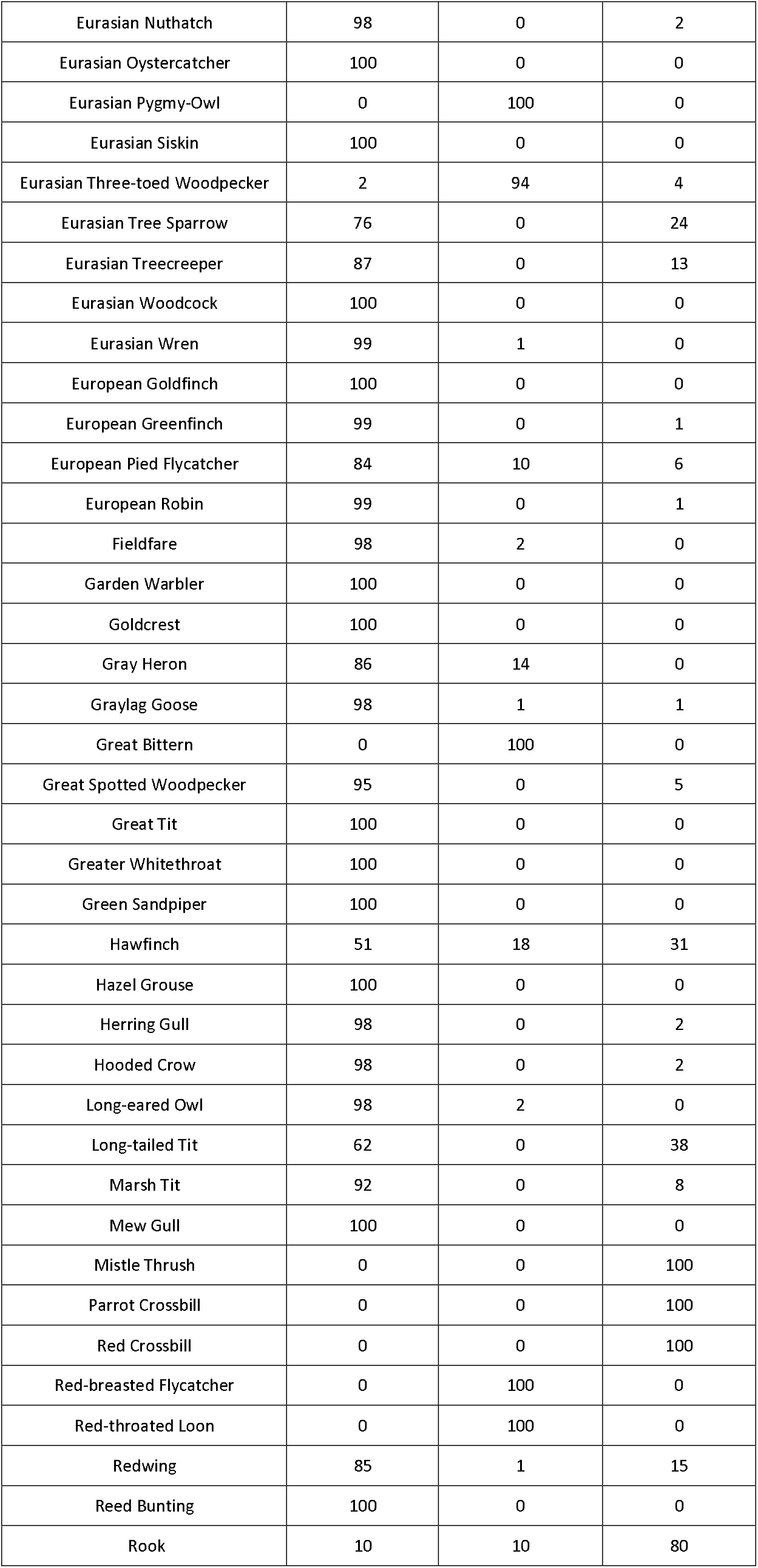
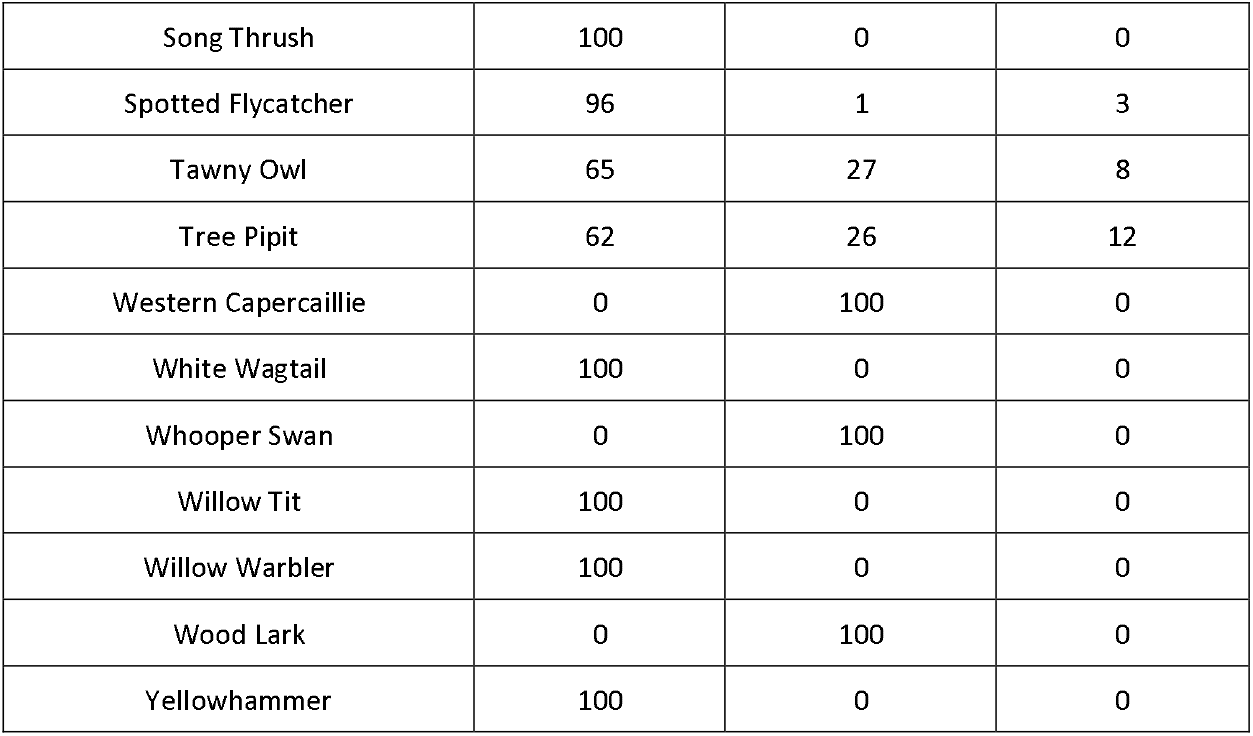
Results of expert ornithologist validation of BirdNET detections. At least 50 random detections, with a model confidence of at least 0.80, were reviewed per species.

**SI Table 4.** Audio Species Distribution Model (aSDM) model hyperparameters, as well as test and train set mean absolute error (MAE) for Willow Warbler and Spotted Flycatcher. Predicted aSDM values range from 0.0 to 1.0. One Random Forest model was trained for each species and hyperparameters were chosen over 1000 iterations of randomized parameters to optimize the test set MAEs.

| Species | Train MAE | Test MAE | Random Forest Parameters |
| --- | --- | --- | --- |
| Willow Warbler<br>(*Phylloscopus trochilus*) | 0.0761 | 0.226 | {'n_estimators': 386, 'min_samples_split': 2, 'min_samples_leaf': 1, 'max_features': 'log2', 'max_depth': 80, 'bootstrap': True} |
| Spotted Flycatcher<br>(*Muscicapa striata*) | 0.0301 | 0.0704 | {'n_estimators': 176, 'min_samples_split': 2, 'min_samples_leaf': 1, 'max_features': None, 'max_depth': 100, 'bootstrap': True} |

## Notes

https://doi.org/10.5281/zenodo.11238950

### Competing Interest Statement

The authors have declared no competing interest.

### Summary of Updates

Additional literature context in introduction and discussion. Cleaning of figures.

## References

1. Saino, N. et al. Climate warming, ecological mismatch at arrival and population decline in migratory birds. Proc. R. Soc. B Biol. Sci. 278, 835–842 (2011).

2. Blicharska, M. et al. Biodiversity’s contributions to sustainable development. Nat. Sustain. 2, 1083–1093 (2019).

3. Robertson, E. P. et al. Decoupling of bird migration from the changing phenology of spring green-up. Proc. Natl. Acad. Sci. 121, e2308433121 (2024).

4. Zwerts, J. A. et al. FSC-certified forest management benefits large mammals compared to non-FSC. Nature https://doi.org/10.1038/s41586-024-07257-8 (2024) doi:10.1038/s41586-024-07257-8.

5. Sauer, J. R. et al. The first 50 years of the North American Breeding Bird Survey. The Condor 119, 576–593 (2017).

6. Linduska, J. P. & Surber, E. Effects of DDT and other insecticides on fish and wildlife: Summary of investigations during 1947. (1948).

7. Hudson, M.-A. R. et al. The role of the North American Breeding Bird Survey in conservation. The Condor 119, 526–545 (2017).

8. J. A. Kålås, I. J. Øien, & B. Stokke. Norwegian breeding bird monitoring scheme - Norsk hekkefuglovervåking. GBIF 10.15468/6jmw2e (2022).

9. La Sorte, F. A. et al. Global change and the distributional dynamics of migratory bird populations wintering in Central America. Glob. Change Biol. 23, 5284–5296 (2017).

10. Zhang, G. Spatial and Temporal Patterns in Volunteer Data Contribution Activities: A Case Study of eBird. ISPRS Int. J. Geo-Inf. 9, 597 (2020).

11. Johnston, A. et al. Analytical guidelines to increase the value of community science data: An example using eBird data to estimate species distributions. Divers. Distrib. 27, 1265–1277 (2021).

12. Horton, K. G. et al. Artificial light at night is a top predictor of bird migration stopover density. Nat. Commun. 14, 7446 (2023).

13. Besson, M. et al. Towards the fully automated monitoring of ecological communities. Ecol. Lett. 25, 2753–2775 (2022).

14. Gibb, R., Browning, E., Glover□Kapfer, P. & Jones, K. E. Emerging opportunities and challenges for passive acoustics in ecological assessment and monitoring. Methods Ecol. Evol. 10, 169–185 (2019).

15. Stowell, D. Computational bioacoustics with deep learning: a review and roadmap. PeerJ 10, e13152 (2022).

16. Pérez□Granados, C. BirdNET: applications, performance, pitfalls and future opportunities. Ibis 165, 1068–1075 (2023).

17. Kahl, S. BirdNET: A deep learning solution for avian diversity monitoring. Ecol. Inform. 10 (2021).

18. Cretois, B., Rosten, C. M. & Sethi, S. S. Voice activity detection in eco□acoustic data enables privacy protection and is a proxy for human disturbance. Methods Ecol. Evol. 13, 2865–2874 (2022).

19. Astaras, C., Linder, J. M., Wrege, P., Orume, R. D. & Macdonald, D. W. Passive acoustic monitoring as a law enforcement tool for Afrotropical rainforests. Front. Ecol. Environ. 15, 233–234 (2017).

20. Prince, P. et al. Deploying Acoustic Detection Algorithms on Low-Cost, Open-Source Acoustic Sensors for Environmental Monitoring. Sensors 19, 553 (2019).

21. Wrege, P. H., Rowland, E. D., Keen, S. & Shiu, Y. Acoustic monitoring for conservation in tropical forests: examples from forest elephants. Methods Ecol. Evol. 8, 1292–1301 (2017).

22. Aniceto, A. S., Ferguson, E. L., Pedersen, G., Tarroux, A. & Primicerio, R. Temporal patterns in the soundscape of a Norwegian gateway to the Arctic. Sci. Rep. 12, 7655 (2022).

23. Sethi, S. S. et al. SAFE Acoustics: A. open□source, real□time eco□acoustic monitoring network in the tropical rainforests of Borneo. Methods Ecol. Evol. 11, 1182–1185 (2020).

24. Sethi, S. S. et al. Automatic vocalisation detection delivers reliable, multi-faceted, and global avian biodiversity monitoring. Preprint at 10.1101/2023.09.14.557670 (2023).

25. Cretois, B. et al. Snowmobile noise alters bird vocalization patterns during winter and pre□breeding season. J. Appl. Ecol. 61, 340–350 (2024).

26. Klingbeil, B. T. & Willig, M. R. Bird biodiversity assessments in temperate forest: the value of point count versus acoustic monitoring protocols. PeerJ 3, e973 (2015).

27. Furnas, B. J. & McGrann, M. C. Using occupancy modeling to monitor dates of peak vocal activity for passerines in California. The Condor 120, 188–200 (2018).

28. Slagsvold, T. Bird Song Activity in Relation to Breeding Cycle, Spring Weather, and Environmental Phenology. Ornis Scand. 8, 197 (1977).

29. Ware, H. E., McClure, C. J. W., Carlisle, J. D. & Barber, J. R. A phantom road experiment reveals traffic noise is an invisible source of habitat degradation. Proc. Natl. Acad. Sci. 112, 12105–12109 (2015).

30. Halfwerk, W., Lohr, B. & Slabbekoorn, H. Impact of Man-Made Sound on Birds and Their Songs. in Effects of Anthropogenic Noise on Animals (eds Slabbekoorn, H., Dooling, R. J., Popper, A. N. & Fay, R. R. vol. 66 209–242 (Springer New York, New York, NY, 2018).

31. Butler, C. J. The disproportionate effect of global warming on the arrival dates of short-distance migratory birds in North America. Ibis 145, 484–495 (2003).

32. Oliver, R. Y. et al. Eavesdropping on the Arctic: Automated bioacoustics reveal dynamics in songbird breeding phenology. Sci. Adv. 4, eaaq1084 (2018).

33. Roe, P. et al. The Australian Acoustic Observatory. Methods Ecol. Evol. 12, 1802–1808 (2021).

34. Desjonquères, C. et al. Acoustic species distribution models (aSDMs): A framework to forecast shifts in calling behaviour under climate change. Methods Ecol. Evol. 13, 2275– 2288 (2022).

35. Wood, C. M. & Kahl, S. Guidelines for appropriate use of BirdNET scores and other detector outputs. J. Ornithol. 10.1007/s10336-024-02144-5 (2024) doi:10.1007/s10336-024-02144-5.

36. Hijmans, R. & University of California, Berkeley, Museum of Vertebrate Zoology. First-level Administrative Divisions, Norway, 2015. UC Berkeley, Museum of Vertebrate Zoology (2015).

37. Sethi, S. S. et al. Characterizing soundscapes across diverse ecosystems using a universal acoustic feature set. Proc. Natl. Acad. Sci. 117, 17049–17055 (2020).

38. Fink, D. et al. eBird Status and Trends, Data Version: 2022. Cornell Lab of Ornithology 10.2173/ebirdst.2022 (2023).

39. Gil, D., Graves, J. A. & Slater, P. J. B. Seasonal patterns of singing in the willow warbler: evidence against the fertility announcement hypothesis. Anim. Behav. 58, 995–1000 (1999).

40. Moran, I. G. et al. Diel and seasonal patterns of variation in the singing behaviour of Savannah Sparrows (Passerculus sandwichensis). Avian Res. 10, 26 (2019).

41. Mauriño, R. A., Zurano, J. P., Zurita, G. A., Di Giacomo, A. S. & De Araújo, C. B. Optimizing passive acoustic monitoring (PAM) for biodiversity studies: Using species–area relationship (SAR) to predict species richness. Ornithol. Appl. duaf063 (2025) doi:10.1093/ornithapp/duaf063.

42. Hedlund, J. S. U., Jakobsson, S., Kullberg, C. & Fransson, T. Long□term phenological shifts and intra□specific differences in migratory change in the willow warbler Phylloscopus trochilus. J. Avian Biol. 46, 97–106 (2015).

43. Clement, P. Willow Warbler (Phylloscopus trochilus). in Birds of the World (eds Billerman, S. M., Keeney, B. K., Rodewald, P. G. & Schulenberg, T. S. (Cornell Lab of Ornithology, 2020). doi:10.2173/bow.wlwwar.01.

44. Mlodinow, S. G., Pyle, P. & Boesman, P. F. D. Common Chiffchaff (Phylloscopus collybita). in Birds of the World (eds Billerman, S. M., Keeney, B. K., Rodewald, P. G. & Schulenberg, T. S. (Cornell Lab of Ornithology, 2025). doi:10.2173/bow.comchi1.02.

45. Taylor, B. Spotted Flycatcher (Muscicapa striata). in Birds of the World (eds Billerman, S. M., Keeney, B. K., Rodewald, P. G. & Schulenberg, T. S. (Cornell Lab of Ornithology, 2020). doi:10.2173/bow.spofly1.01.

46. Barrett, R. T. The dependence of long-distance migration to North Norway on environmental conditions in the wintering area and en route. Ornis Nor. 40, 14 (2017).

47. Burger, C. et al. Climate change, breeding date and nestling diet: how temperature differentially affects seasonal changes in pied flycatcher diet depending on habitat variation. J. Anim. Ecol. 81, 926–936 (2012).

48. Lundberg, A., Alatalo, R. V., Carlson, A. & Ulfstrand, S. Biometry, Habitat Distribution and Breeding Success in the Pied Flycatcher Ficedula hypoleuca. Ornis Scand. 12, 68 (1981).

49. Hedenström, A. & Pettersson, J. Migration routes and wintering areas of Willow Warblers Phylloscopus trochilus (L.) ringed in Fennoscandia. Ornis Fenn. 64, (1987).

50. Yohannes, E., Biebach, H., Nikolaus, G. & Pearson, D. J. Migration speeds among eleven species of long□distance migrating passerines across Europe, the desert and eastern Africa. J. Avian Biol. 40, 126–134 (2009).

51. Lyapustin, A. MODIS/Terra+Aqua Vegetation Index from MAIAC, Daily L3 Global 0.05Deg CMG V061. NASA EOSDIS Land Processes Distributed Active Archive Center 10.5067/MODIS/MCD19A3CMG.061 (2023).

52. Barrett, R. T. The phenology of spring bird migration to north Norway. Bird Study 49, 270–277 (2002).

53. Lameris, T. K. et al. Arctic Geese Tune Migration to a Warming Climate but Still Suffer from a Phenological Mismatch. Curr. Biol. 28, 2467-2473.e4 (2018).

54. Newton, I. Weather□related mass□mortality events in migrants. Ibis 149, 453–467 (2007).

55. Breiman, L. Random Forests. Mach. Learn. 45, 5–32 (2001).

56. Cristian Lussana. seNorge/TX: daily maximum temperature over Norway (22.09) [Data Set]. Zenodo 10.5281/zenodo.6965960 (2022).

57. Cristian Lussana. seNorge/TN: daily minimum temperature over Norway (22.09) [Data set]. Zenodo 10.5281/zenodo.6965929 (2022).

58. Cristian Lussana. seNorge/TG: daily mean temperature over Norway (22.09) [Data set]. Zenodo 10.5281/zenodo.6965866 (2022).

59. Cristian Lussana. seNorge/RR: daily total precipitation amounts over Norway (22.09) [Data set. Zenodo 10.5281/zenodo.6965906 (2022).

60. Kartverket (Norwegian Geographical Survey). Norway National Detailed Elevation Model. Kartverket.

61. Copernicus Climate Change Service, Climate Data Store. Land cover classification gridded maps from 1992 to present derived from satellite observations. Copernicus Climate Change Service (C3S) Climate Data Store (CDS) 10.24381/cds.006f2c9a (2019).

62. Bota, G., Traba, J., Sardà-Palomera, F., Giralt, D. & Pérez-Granados, C. Acoustic Monitoring of Diurnally Migrating European Bee-Eaters Agrees with Data Derived from Citizen Science. Ardea 108, (2020).

63. Zamora-Marín, J. M., Zamora-López, A., Calvo, J. F. & Oliva-Paterna, F. J. Comparing detectability patterns of bird species using multi-method occupancy modelling. Sci. Rep. 11, 2558 (2021).

64. Knight, E. et al. Individual identification in acoustic recordings. Trends Ecol. Evol. 39, 947– 960 (2024).

65. Heath, B. E. et al. Spatial ecosystem monitoring with a Multichannel Acoustic Autonomous Recording Unit (MAARU). Methods Ecol. Evol. 15, 1568–1579 (2024).

66. Stowell, D., Petrusková, T., Šálek, M. & Linhart, P. Automatic acoustic identification of individuals in multiple species: improving identification across recording conditions. J. R. Soc. Interface 16, 20180940 (2019).

67. Waddell, E. E., Rasmussen, J. H. & Širović, A. Applying Artificial Intelligence Methods to Detect and Classify Fish Calls from the Northern Gulf of Mexico. J. Mar. Sci. Eng. 9, 1128 (2021).

68. Cretois, B. et al. TABMON – real-time acoustic biodiversity monitoring across Europe. Preprint at 10.32942/X2236J (2026).

69. Van Ommen Kloeke, E. et al. ARISE: Building an infrastructure for species recognition and biodiversity monitoring in the Netherlands. Biodivers. Inf. Sci. Stand. 6, e93613 (2022).

70. Rich, L. N., Furnas, B. J., Newton, D. S. & Brashares, J. S. Acoustic and camera surveys inform models of current and future vertebrate distributions in a changing desert ecosystem. Divers. Distrib. 25, 1441–1456 (2019).

71. Sethi, S. S., Kovac, M., Wiesemüller, F., Miriyev, A. & Boutry, C. M. Biodegradable sensors are ready to transform autonomous ecological monitoring. Nat. Ecol. Evol. 6, 1245–1247 (2022).

72. Fitzpatrick, J. W., Gill, F., Powers, M., Wells, J. V. & Rosenberg, K. V. Introducing eBird: The Union of Passion and Purpose. North Am. Birds 56, (2002).

73. OpenStreetMaps Contributors. OpenStreetMaps Norway Dataset. Geofabrik (2022).

74. Sethi, S. https://github.com/bugg-resources. Bugg (2022).

75. Heath, B. E., Sethi, S. S., Orme, C. D. L., Ewers, R. M. & Picinali, L. How index selection, compression, and recording schedule impact the description of ecological soundscapes. Ecol. Evol. 11, 13206–13217 (2021).

76. K. Lisa Yang Center for Conservation Bioacoustics. BirdNET-lite. (2022).

77. Beery, S., Cole, E., Parker, J., Perona, P. & Winner, K. Species Distribution Modeling for Machine Learning Practitioners: A Review. in ACM SIGCAS Conference on Computing and Sustainable Societies (COMPASS) 329–348 (ACM, Virtual Event Australia, 2021). doi:10.1145/3460112.3471966.

